# *Parkia javanica* extracts exhibits tissue regeneration potential in Zebrafish (*Danio rerio*)

**DOI:** 10.1101/2025.10.12.681930

**Authors:** Antara Bhuyan, Achinta Singha, Krithika Kalladka, Rajeshwari Vittal, Partha Saha, Samir Kumar Sil, Anirban Chakraborty, Gunimala Chakraborty

## Abstract

*Parkia javanica* is a medicinal plant acknowledged for its diverse pharmacological features, but its biological effects, like regeneration and wound-healing properties, in the zebrafish animal model (*Danio rerio*) is unexplored. The purpose of this study was to determine the caudal fin tissue regeneration and antioxidant potential in response to *Parkia javanica* fruit and bark extracts on *Danio rerio*. The *Danio rerio* caudal fin was amputated and subsequently was treated with *Parkia javanica* fruit and bark extracts at 0.346µg/mL and 2.86µg/mL respectively. The regenerative effects of *Parkia javanica* fruit and bark extracts were evaluated through morphological analysis and dorso-ventral patterning. Additionally, the antioxidant properties of *Parkia javanica* fruit and bark extracts, along with the mechanistic insights, were evaluated using qRT-PCR. We found that both the *Parkia javanica* fruit and bark extracts displayed substantial antioxidant capacity with upregulation of key genes like *Cat* and *Sod1*. Further, the extracts demonstrated significant fin regeneration compared to the control group. We observed that both the *Parkia javanica* fruit and bark extracts possess tissue regeneration properties by upregulating key genes, like *Anxa2a, Anxa2b*, and *Wnt3a*. All these findings provide novel insights into the molecular mechanisms underlying the tissue repair and regeneration effects of *Parkia javanica* fruit and bark extracts and may pave the way for the development of novel regenerative therapeutic strategies.

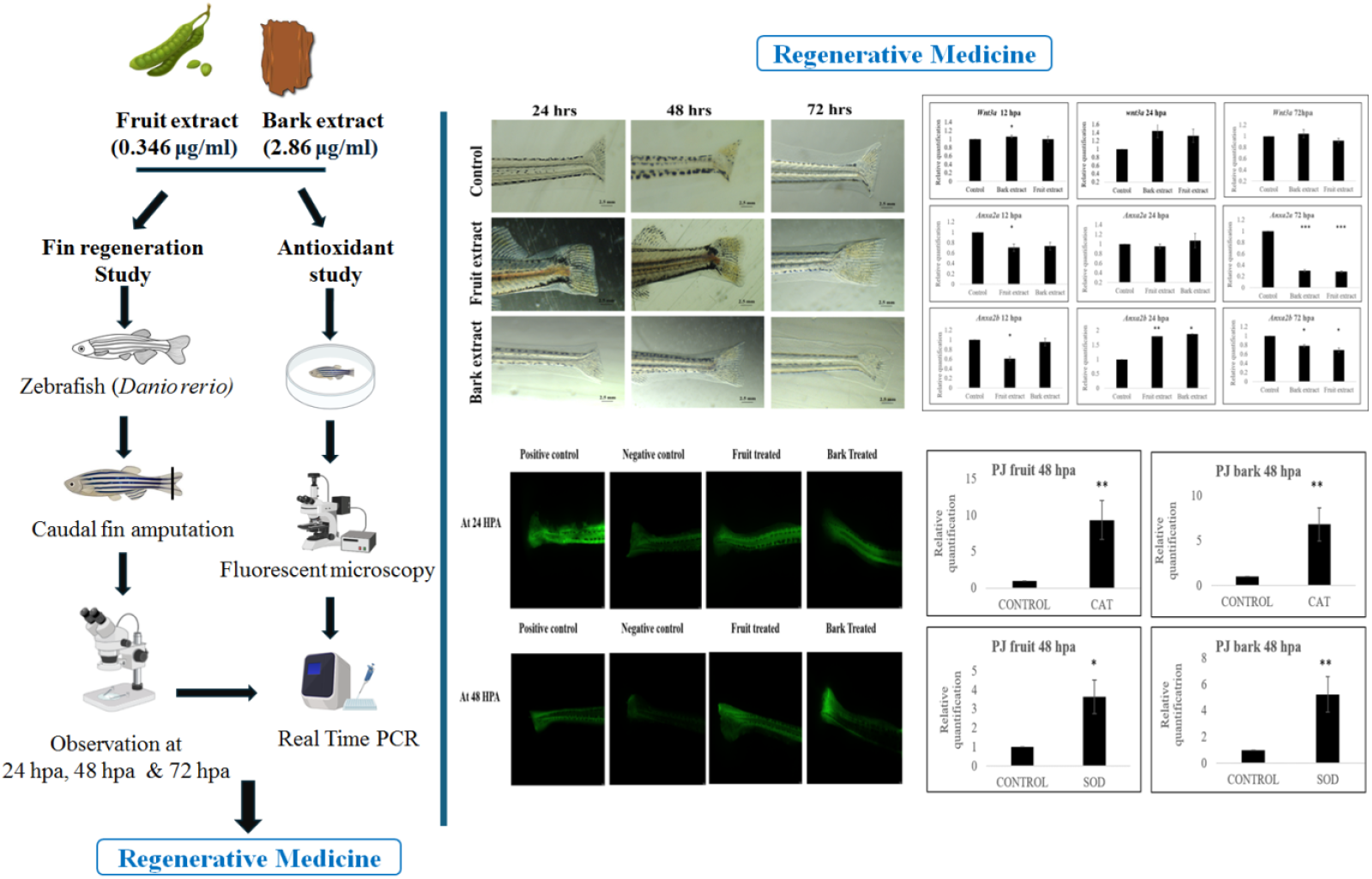

## Introduction

Regenerative medicine aims to restore damaged tissues and organs. Traditional bioactive compounds, such as growth factors and cytokines, have shown considerable promise in tissue regeneration, but they have significant drawbacks, such as limited availability, high cost, short half-life, and side effects [1]. In recent years, plant-derived bioactive compounds have drawn more attention due to their accessibility, cost-effectiveness, and safety, as well as their significant therapeutic advantages, including anti-inflammatory, antibacterial, antioxidative, and pro-angiogenic properties, along with the potential to induce stem cell differentiation [1–2]. The use of these medicinal plants is beneficial for health, and their therapeutic purposes have increased during the last five years [3–4]. Indeed, several medical plant extracts are now recommended in many developed nations, including the United Kingdom, Germany, and France [5–6]. Many medicinal plant extracts have been employed to enhance stem cell proliferation and differentiation, as well as to stimulate tissue regeneration, resulting in the rehabilitation of damaged or diseased tissues [7–8]. A number of natural bioactive substances are being utilized globally to treat and prevent different diseases due to their multi-target, multi-layer, and multi-pathway properties [9]. *Parkia* is a genus of flowering plants belonging to the family Fabaceae that comprises about 34 species across all tropical countries and is considered as food and folklore medicine. The tribal people of north-east India traditionally use this plant, especially *Parkia javanica* to cure various ailments, including anticancer, antibacterial, antihypertensive, antiulcer, antidiabetic, anti-inflammatory, antioxidant, antimalarial, hepatoprotective, antidiarrheal, and wound healing properties [10–11]. Various other plants like *Pistacia atlantica, Zataria multiflora, Trifolium repens, Quercus infectoria*, and *Salvia officinalis* also demonstrated the wound-healing properties both in *in-vitro* and *in-vivo* wound models [12–16]. Studies also demonstrated that blueberry and catechin had a dose-dependent activity on human bone marrow proliferation [17–18]. However, there is no scientific evidence supporting *Parkia javanica*’s wound healing properties in zebrafish animal model. The zebrafish has emerged as a valuable and cost-effective animal model to explore tissue regeneration due to its ability to regenerate many tissues and organs over short periods of time [19]. Zebrafish (*Danio rerio*) is an excellent vertebrate animal model for studying regeneration because they can regenerate organs/appendages like fins [20–21]. The benefit of studying fin regeneration in zebrafish is that the fins are easily accessible, and the amputation has no significant adverse effect on the animal, and also the regenerated fins can be visible with the necked eyes [22]. Zebrafish, as an animal model, are a useful tool in extending our understanding of pharmacological research toward safer and more effective therapies [23]. A number of research groups have investigated the molecular mechanisms that regulate blastema development and proliferation [24–28]. In our previous investigation, we reported the LC50 of *Parkia javanica* fruit and bark extracts (346.6 mg/L and 28.66 mg/L, respectively), as well as their anti-angiogenic and anti-proliferative effects [29]. The investigation provides us the safe dose of *Parkia javanica* fruit and bark extracts, and considering the evidence, the current study was designed to evaluate the antioxidant and regeneration properties of *Parkia javanica* fruit and bark extracts as a first attempt toward scientifically establishing their role that could be translated into an alternative to regenerative therapy.

## Materials and Methods

### Zebrafish care

The zebrafish were raised in a recirculating system that constantly aerates, ensuring a 14/10 hrs day/night cycle with a fixed temperature of 28.5°C. The breeding was set up late in the evening, and spawning began shortly after light appeared. The eggs were collected, washed, and placed in a fresh petri dish with E3 embryonic rearing media. They were housed at a 28°C-cooling incubator (Panasonic, Japan). For this investigation, healthy fish of similar ages were selected [30].

### Evaluation of caudal fin regeneration

In short, the juvenile AB wild-type strain zebrafish was used for this experiment. Prior to the amputation, the fish was anesthetized with tricaine (0.02%), and the caudal fin was amputated at identical distances with sterilized surgical blades. The amputated fish were carefully transferred in separate tanks, and the growth of the fin was observed under standard laboratory conditions. The zebrafish were exposed to *Parkia javanica* fruit and bark extracts at concentrations of 0.346 µg/mL (1/1000^th^ of LC50) and 2.86 µg/mL (1/10^th^ of LC50) respectively, and the control group received only the E3 media. The optimum dose was calculated based on the previous results reported from our lab. The caudal fin regeneration was assessed, and the tail regrowth pattern was examined at several time points (24 hpa, 48 hpa, and 72 hpa) [31]. All the images were acquired at each time point using a stereo microscope (Leica S9D, Japan). The experiment was repeated three times, and the average mean value was calculated.

### Reactive oxygen species or ROS measurement

Post-amputation oxidative stress results in the production of reactive oxygen species (ROS). To investigate the antioxidant potential of *Parkia javanica* fruit and bark extracts, zebrafish embryos were administered 2′,7′-dihydrodichlorofluorescein diacetate (H2DCFDA), which was oxidized by ROS to produce a green fluorescent dichlorofluorescein (DCF) compound. The zebrafish were treated to *Parkia javanica* fruit and bark extracts at 0.346 µg/mL (1/1000^th^ of LC50) and 2.86 µg/mL (1/10^th^ of LC50), and ROS generation was measured in response to amputation at 48 hpa [32]. Following incubation, the fish were rinsed with 1X PBS, and the images were acquired using a fluorescent microscope (Leica-DFC7000 LED). The experiment was carried out three times independently.

### Quantitative real-time PCR (qPCR)

Quantitative real-time PCR was performed to analyse the expression of *Anxa2a, Anxa2b, and Wnt3a* genes in order to identify the key mediators of dorso-ventral patterning and the regulation of growth control during fin regeneration. We also explore the key genes associated with oxidative stress, like the *Cat* and *Sod1* genes. In short, for quantitative analysis of dorso-ventral patterning and for oxidative stress, the total RNA was extracted from the fins using the TRIzol method, and the RNA concentration and purity were checked using Multiscan Sky spectrophotometer (model), followed by cDNA synthesis using PrimeScript RT reagent kit (Cat. # RR037A) according to the manufacturer’s protocol. The quantitative real-time PCR was performed using TaKaRa bio kit, Japan and the master mix composition used for real-time PCR is given in Table 1.

**Table 1.** Master mix for Real Time.

| Material | Volume |
| --- | --- |
| SYBR Green Supermix | 5µl |
| Forward primer | 0.4µl |
| Reverse Primer | 0.4µl |
| ROX Dye II | 0.2µl |
| cDNA | 2µl |
| De-ionised water | Made up to 10µl |

The cycling conditions included an initial denaturation at 95°C for 2 minutes, followed by 95°C for 15 seconds, annealing at 56°C-60°C for 30 seconds, and extension at 72°C for 1 minute, repeated for 40 cycles with the default melt curve at 95°C for 15 seconds, 60°C for 1 minute, and 95°C for 15 seconds. The p-value was calculated using a t-test, and a value of <0.05 was considered statistically significant. Quantitative analysis of Wnt3a, Anxa2a, and Anxa2b was repeated three times for each time point, and results were averaged. The following primers were used for qRT-PCR (Table 2):

**Table 2.**
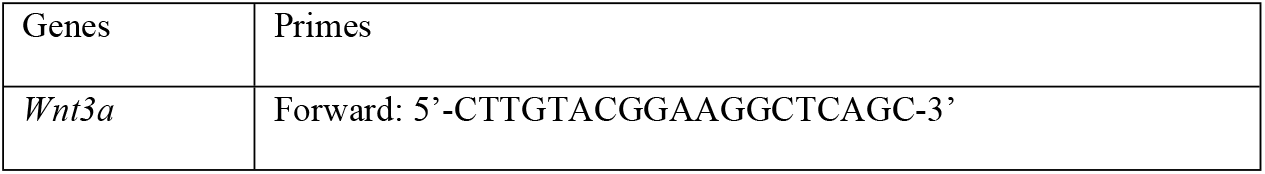
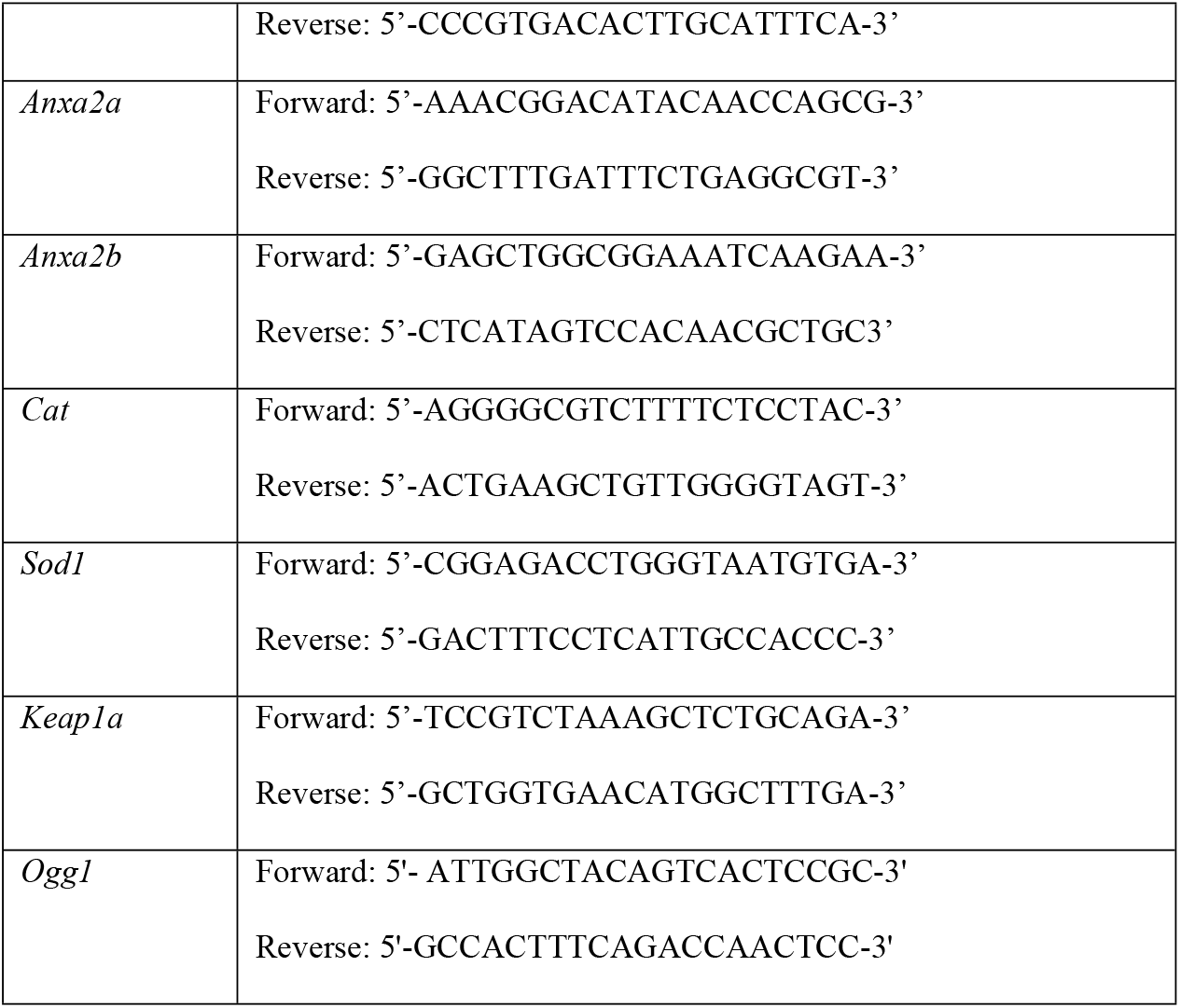
Sequences of primers used for qRT-PCR.

### Statistical analysis

All the data were expressed as the mean ± standard deviation from three independent experiments. One-way ANOVA was used to compare distinct groups, followed by paired t-tests. A p-value < 0.05 indicated a significant difference between groups.

## Results

### *Parkia javanica* extracts promoted zebrafish fin regeneration

To study the potential impact of *Parkia javanica* fruit and bark extracts, we amputated the caudal fin of zebrafish and exposed them to *Parkia javanica* fruit and bark extracts at 0.346 µg/mL (1/1000^th^ of LC50) and 2.86 µg/mL (1/10^th^ of LC50), respectively. The length of the regenerated fin was measured at 24, 48, and 72 dpa. The result demonstrated that zebrafish embryos exposed to 0.346 µg/mL and 2.86 µg/mL *Parkia javanica* fruit and bark extracts had showed higher fin regeneration compared to the control group. We also observed that the fruit extract shown higher fin regeneration compared to bark extract (Figure 1). These findings indicated that *Parkia javanica* fruit and bark extracts increase zebrafish fin regeneration that could be translated into an alternative to regenerative therapeutics.

**Figure 1.**
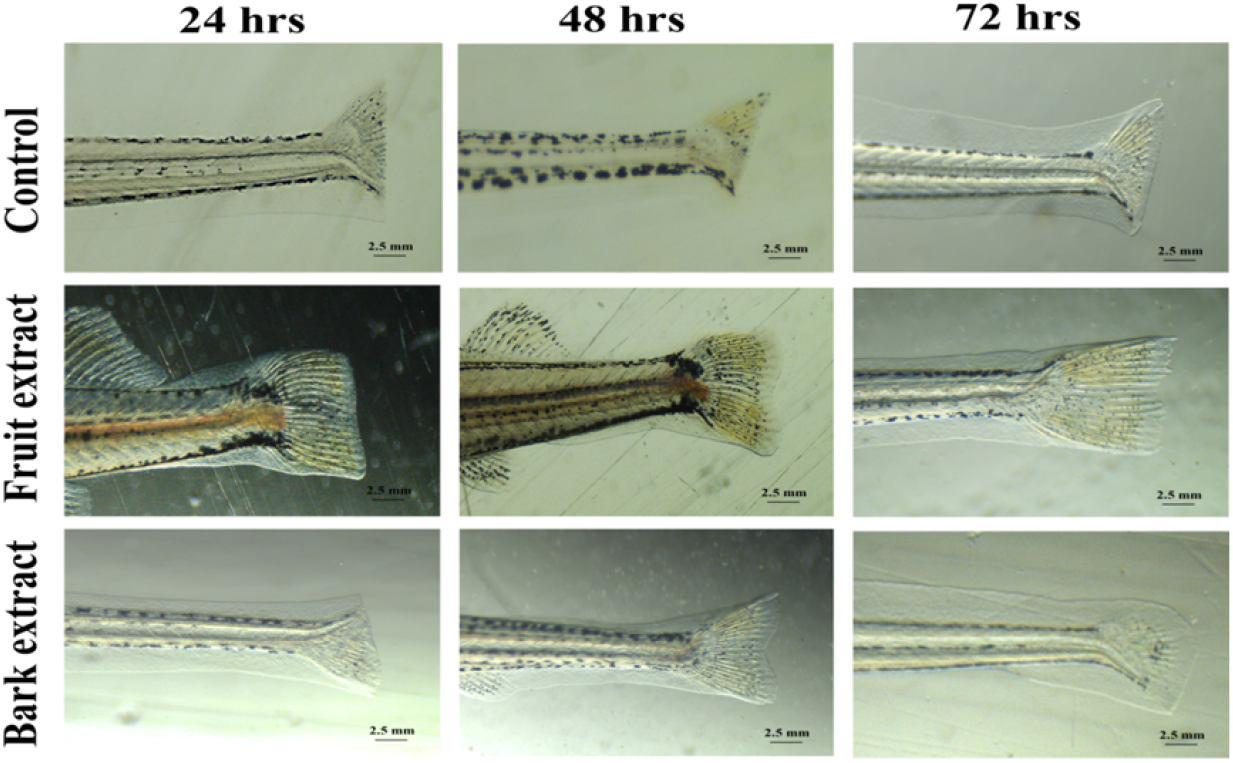
Assessment of *Parkia javanica* fruit and bark extracts caudal fin regeneration. The amputated juvenile zebrafish were transferred to E3 media without drug (control group) and E3 media with drugs (treated groups) for 24 hpa, 48 hpa, and 72 hpa. The E3 media and drugs were replenished daily after taking the images at regular time intervals. The lengths of each regenerated fish in each group were measured at a scale bar of 2.5 mm, and the experiment was repeated three times.

Reduction of reactive oxygen species (ROS) upon exposure to *Parkia javanica* fruit and bark extracts The antioxidant effect of *Parkia javanica* fruit and bark extracts was determined by quantifying the ROS levels using the 2′,7′-dichlorodihydrofluorescein diacetate (H2DCFDA) assay. Our data showed the relative fluorescence intensity was higher in the control group compared to that in larvae exposed to *Parkia javanica* fruit and bark extracts (Figure 2). The result confirmed that the *Parkia javanica* fruit and bark extracts have potent antioxidant activity by reducing the reactive oxygen species. We found that the fruit extract showed lower fluorescent intensity compared to the bark extract.

**Figure 2.**
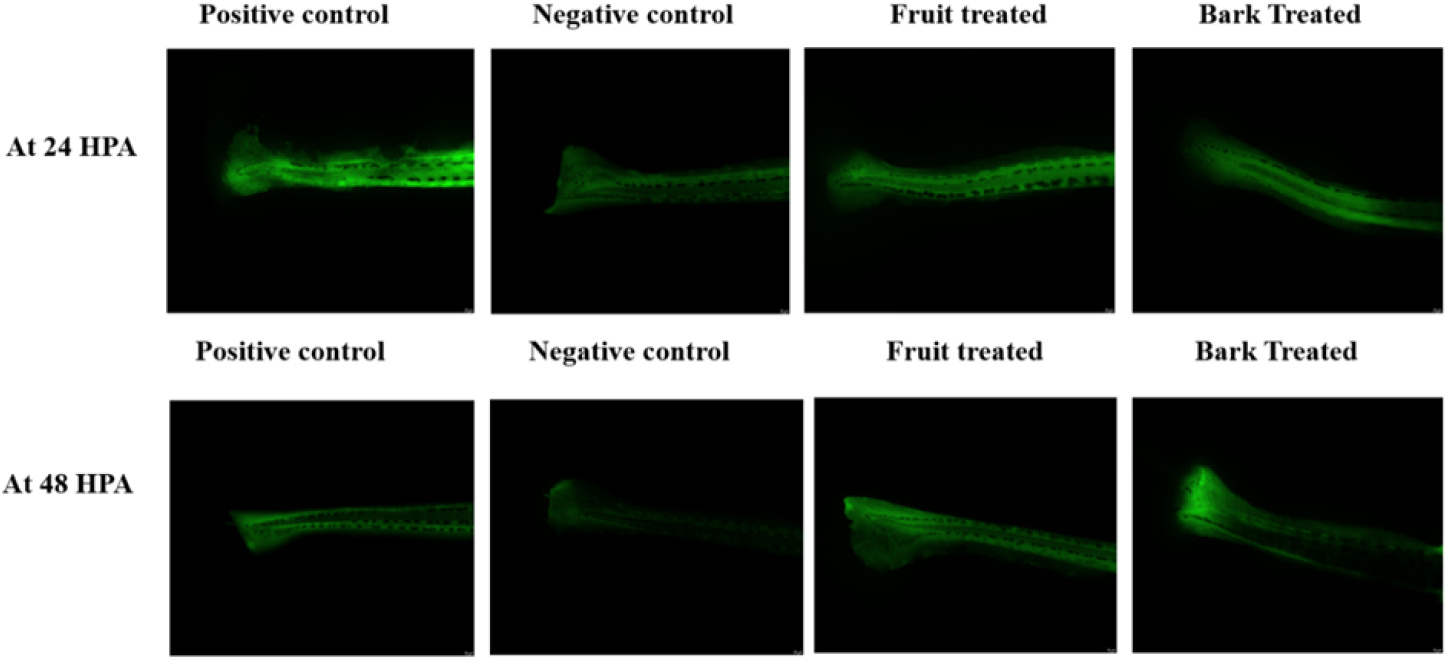
Investigation of antioxidant potential of *Parkia javanica* fruit and bark extracts on a zebrafish animal model. The images represent the fluorescence intensity of the tail region in the control group and treated groups (fruit and bark extract) after the exposure.

### Effect of *Parkia javanica* fruit and bark extract on the expression of oxidative stress response genes *Cat* and *Sod1*

The expression levels of Cat and Sod1 were determined by using RT-qPCR, and we observed that the treatment groups had considerably higher expression levels of Cat and Sod1 genes than the control group (Figure 3). We found that the Cat gene was upregulated by around 8-fold in the *Parkia javanica* fruit extract-treated group, whereas the *Parkia javanica* bark extract showed 6-fold. The SOD gene was also upregulated by 3.5-fold in the fruit and 5-fold in the bark extract-treated groups.

**Figure 3.**
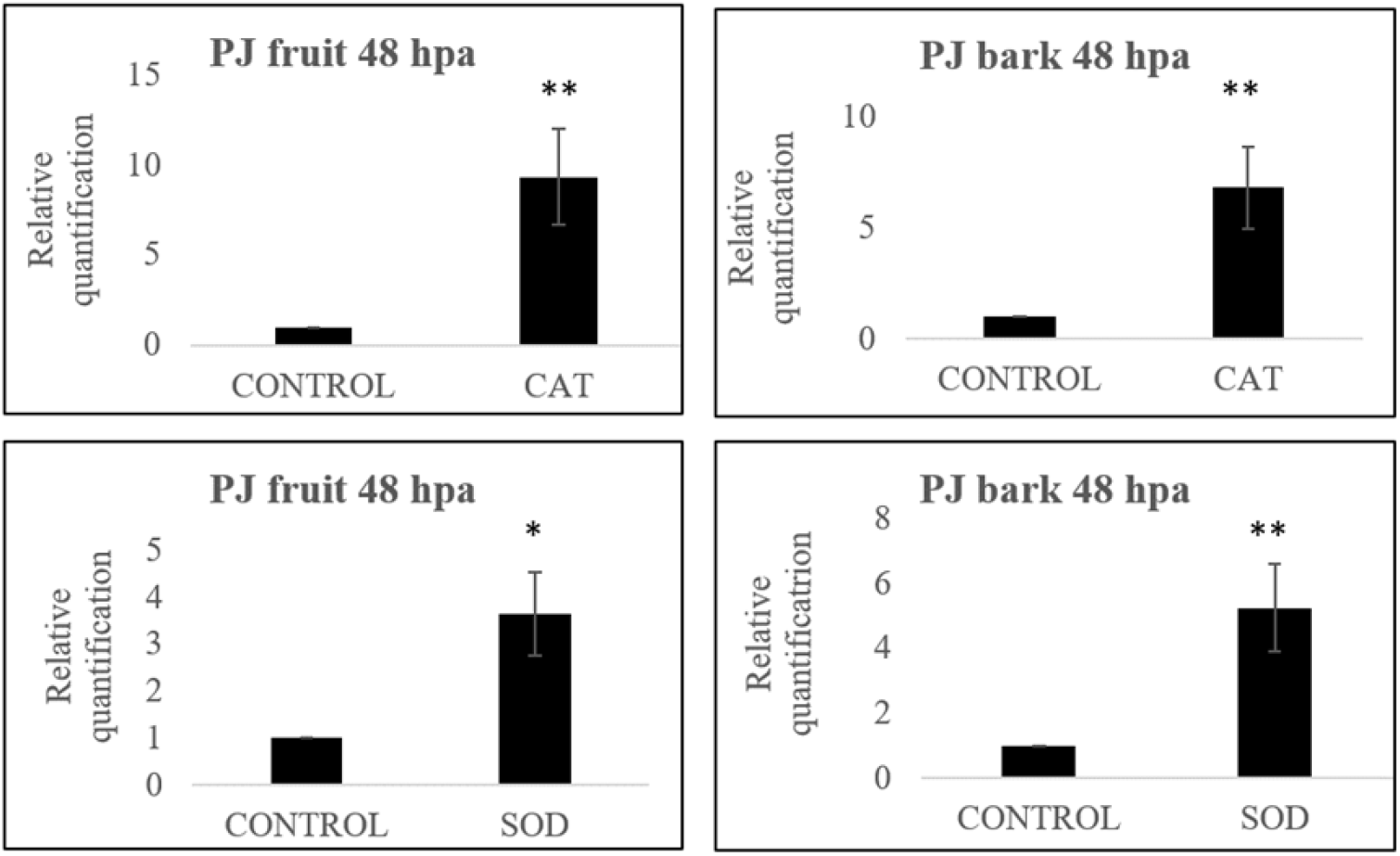
The relative expression of oxidative stress response genes in zebrafish larvae treated with *Parkia javanica* fruit and bark extract at different time points. The bar diagrams show the expression of Cat and Sod, and the data are represented as mean ± SD from three independent replicates. * p< 0.05, ** p< 0.01, *** p< as compared to the control group and *Parkia javanica* fruit and bark extract treatment group.

**Figure 4.**
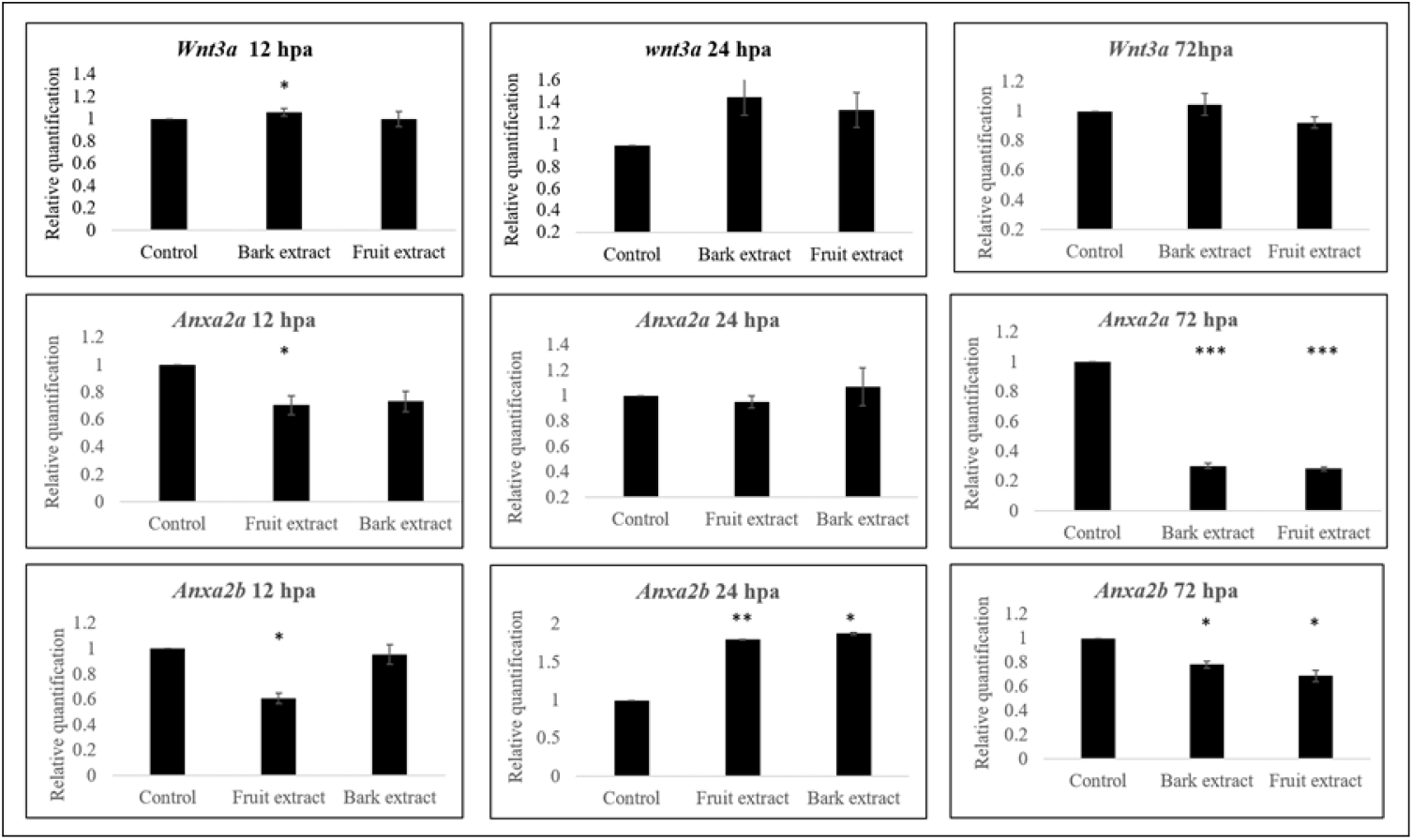
The quantitative real-time PCR analysis of the effects of *Parkia javanica* fruit and bark extract on the zebrafish animal model. The bar diagram depicts the expression of the Anxa2a, Anxa2b, and Wnt3a genes at 24 hpa, 48 hpa, and 72 hpa. The experiment was carried out in triplicate, and the data are represented as mean ± SD from three independent replicates.

### Effect of *Parkia javanica* fruit and bark extract on expression of *Anxa2a, Anxa2b, and Wnt3a*

Quantitative real-time PCR was used to assess the expression of the genes Anxa2a, Anxa2b, and Wnt3a to identify the key mediators of dorso-ventral patterning and understand the importance of these regulator genes during tail fin regeneration. We found that at 24 hpa the expression of the Wnt3a gene was upregulated after the exposure to *Parkia javanica* fruit and bark extracts. Further, we also found that at 72 hpa, the expression of Wnt3a was down, and similar findings were observed for the genes Anxa2a and Anxa2b as well.

## Discussion

The present morphological analysis of the amputated caudal fin of zebrafish demonstrated that exposure to *Parkia javanica* fruit and bark extracts significantly accelerated zebrafish caudal fin regrowth. Under stereomicroscopic observation, treated groups displayed a more rapid closure of the wound; it might be because the *Parkia javanica* extracts potentially include amides, alkaloids, flavonoids, phenolic compounds, terpenoids, quinones, saponins and tannins [33–34] that may act to stimulate cellular proliferation and extracellular matrix deposition within the regenerative blastema. Previous studies have established zebrafish caudal fin regeneration as a multi-phase process involving inflammation, wound healing, and blastema formation [35]. Zebrafish caudal fin tissue regenerates quickly after the amputation, and understanding the differential expression of regeneration genes/proteins provides detailed knowledge of the molecular mechanisms underlying wound healing, proliferation, differentiation, dedifferentiation, and structural regeneration. The increased regeneration observed in the *Parkia javanica* fruit and bark extract-treated groups could be attributed to the presence of bioactive phytochemicals that modulate the signalling pathways, either through direct activation of regenerative cascades or via antioxidant effects that develop a favourable microenvironment for regrowth.

Zebrafish (*Danio rerio*) are a widely used model organism in regenerative biology due to their remarkable ability to regenerate complex tissues, including fins, heart, spinal cord, retina, and even parts of the brain [21, 36]. Tissue injury and regeneration are closely associated with the generation of oxidative stress [37], and upon injury, there is a rapid accumulation of reactive oxygen species (ROS) that impair the regenerative process. To restore the reduction-oxidation balance, organisms activate endogenous antioxidant defense systems, of which superoxide dismutase (SOD) and catalase (CAT) are two pivotal enzymes [38]. These endogenous antioxidants accelerate the wound healing process by neutralizing these free radicals [39]. Activities of these enzymes in response to *Parkia javanica* extracts were analyzed on the zebrafish model organism, and we found that the expression of the SOD gene was upregulated by 3.5-fold in the fruit and 5-fold in the bark extract-treated groups. The coordinated upregulation of SOD and CAT after the injury helps repair the tissue damage from oxidative stress and is also essential for cell proliferation, matrix remodeling, and angiogenesis, all of which are critical for tissue regeneration. Many studies have demonstrated that exposing zebrafish embryos to the plant extract increased the activity of *CAT* and *SOD* that protects against oxidative stress [40–41]. Furthermore, our recent findings confirmed that *Parkia javanica* fruit and bark extracts have great antioxidant potential, performing by directly scavenging free radicals, increasing the cellular antioxidant defense system, and improving the wound healing process.

Tissue regeneration is a tightly regulated biological process involving cell proliferation, migration, differentiation, and extracellular matrix remodeling [42]. These events are regulated by several molecular pathways, including those governed by annexins and Wnt signaling proteins. Among the key regulators in this context are Annexin A2a (Anxa2a), Annexin A2b (Anxa2b), and Wingless-related integration site 3A (Wnt3a), all of which play crucial roles in regulating regeneration in vertebrate models such as zebrafish [43–44]. Together, Anxa2a, Anxa2b, and Wnt3a represent key molecular switches that initiate and regulate the regenerative cascade in response to injury. Their coordinated expression reflects a conserved mechanism of regeneration, making them reliable biomarkers for evaluating the regenerative potential of therapeutic agents. In this work, *Parkia javanica* fruit and bark were employed to investigate tissue regeneration using zebrafish. We observed that at 24 hpa the expression of the Wnt3a gene was upregulated, whereas the expression of Wnt3a was down at 72 hpa, and similar findings were observed for the genes Anxa2a and Anxa2b as well. All these findings highlight the importance of managing oxidative stress; its association with tissue regeneration might be considered a promising approach.

In the present study, we explored whether *Parkia javanica* fruit and bark extract had any regenerative effects on the model system or not. First, we established the reconstruction of fin morphology and observed each fin’s development by comparing fin ray growth to the control group. We also found that *Parkia javanica* fruit and bark extract had shown significant antioxidant and tissue regeneration activity by up-regulating key regulating genes involved in the pathway.

## Conclusion

The results suggest that both fruit and bark extracts may influence key regenerative processes in zebrafish, such as anti-oxidation, cell proliferation, and tissue differentiation at the site of injury. These effects could be attributed to the bioactive compounds present in *Parkia javanica*, which may modulate inflammation, enhance stem cell activation, and support the overall regenerative environment. By stimulating fin regeneration in zebrafish, *Parkia javanica* extracts showed promise as natural agents that could potentially be explored for applications in regenerative medicine, offering insights into new avenues for tissue healing and repair in both aquatic and potentially human systems. However, further studies are necessary to fully understand the mechanisms of action and long-term effects of these extracts to assess their full therapeutic potential.

## Acknowledgement

The infrastructure support provided by Nitte (Deemed to be a University) and Nitte-GOK CoE, AQUAMARIN Government of Karnataka, is gratefully acknowledged.

## Abbreviations

LC50: Lethal Concentration 50%; mg/L, stands for milligrams per liter;
ROS: reactive oxygen species;1
H2DCFDA: 2′,7′-dihydrodichlorofluorescein diacetate;
DCF: dichlorofluorescein;
Anxa2a: Annexin 2a,Anxa2b, 2b
Wnt3a: Wnt family member 3A
CAT: catalase
SOD1: superoxide dismutase 1;
dpa: days post-amputation;
hpa: hours post-amputation;
O_2_^−^: superoxide radicals.

## Funding

None

## Data Availability

All the data are available upon request.

## Conflict of interest

None of the authors has any financial or personal relationships with other people or organizations that could inappropriately influence this work.

## Author contribution

Antara Bhuyan: Original Draft, Methodology, Investigation, Formal Analysis, Data curation, Achinta Singha: Original Draft, Formal analysis, Conceptualization, Krithika Kalladka: Review & Editing, Methodology, Investigation, Formal Analysis, Data curation, Rajeshwari Vittal: Formal analysis, Review & Editing, Partha Saha: Formal analysis, Review & Editing, Samir Kumar Sil: Formal analysis, Review & Editing, Anirban Chakraborty: Review & Editing, Validation, Supervision, Conceptualization, Gunimala Chakraborty: Review & Editing, Visualization, Validation, Supervision, Conceptualization.

## Ethical approval

All zebrafish experiments followed proper national and international standards. In this investigation we used AB wild-type strain adult and juvenile zebrafish, and they were anesthetized before the amputation. All mentioned zebrafish experimental methods were designed and executed in accordance with the animal ethical rules established by the institute with permission reference number NGSMIPS/IAEC/JAN-2023/344.

## Notes

### Competing Interest Statement

The authors have declared no competing interest.

## References

1) Zhou G, Xu R, Groth T, Wang Y, Yuan X, Ye H, Dou X. The combination of bioactive herbal compounds with biomaterials for regenerative medicine. Tissue Engineering Part B: Reviews. 2024 Dec 1;30(6):607–30. 10.1089/ten.teb.2024.0002

2) Valentino A, Di Cristo F, Bosetti M, Amaghnouje A, Bousta D, Conte R, Calarco A. Bioactivity and delivery strategies of phytochemical compounds in bone tissue regeneration. Applied Sciences. 2021 May 31;11(11):5122. 10.3390/app11115122

3) Banjari I, Misir A, Šavikin K, Jokić S, Molnar M, De Zoysa HK, Waisundara VY. Antidiabetic effects of Aronia melanocarpa and its other therapeutic properties. Frontiers in Nutrition. 2017 Nov 6;4:53. 10.3389/fnut.2017.00053

4) Yatoo MI, Dimri U, Gopalakrishnan A, Karthik K, Gopi M, Khandia R, Saminathan M, Saxena A, Alagawany M, Farag MR, Munjal A. Beneficial health applications and medicinal values of Pedicularis plants: A review. Biomedicine & Pharmacotherapy. 2017 Nov 1;95:1301–13. 10.1016/j.biopha.2017.09.041

5) Thomford NE, Senthebane DA, Rowe A, Munro D, Seele P, Maroyi A, Dzobo K. Natural Products for Drug Discovery in the 21st Century: Innovations for Novel Drug Discovery. Int J Mol Sci. 2018 May 25;19(6):1578. doi:10.3390/ijms19061578

6) Woo CS, Lau JS, El-Nezami H. Herbal medicine: toxicity and recent trends in assessing their potential toxic effects. InAdvances in botanical research 2012 Jan 1 (Vol. 62, pp. 365–384). Academic Press. 10.1016/B978-0-12-394591-4.00009-X

7) Amri B, Martino E, Vitulo F, Corana F, Ben-Kaâb LB, Rui M, Rossi D, Mori M, Rossi S, Collina S. Marrubium vulgare L. leave extract: Phytochemical composition, antioxidant and wound healing properties. Molecules. 2017 Oct 28;22(11):1851. 10.3390/molecules22111851

8) Nimma VL, Talla HV, Bairi JK, Gopaldas M, Bathula H, Vangdoth S. Holistic healing through herbs: Effectiveness of Aloe vera on post extraction socket healing. Journal of clinical and diagnostic research: JCDR. 2017 Mar 1;11(3):ZC83. doi:10.7860/JCDR/2017/21331.9627

9) Huang J, Huang N, Mao Q, Shi J, Qiu Y. Natural bioactive compounds in Alzheimer’s disease: from the perspective of type 3 diabetes mellitus. Frontiers in Aging Neuroscience. 2023 Mar 16;15:1130253. 10.3389/fnagi.2023.1130253

10) Saleh MS, Jalil J, Zainalabidin S, Asmadi AY, Mustafa NH, Kamisah Y. Genus Parkia: Phytochemical, medicinal uses, and pharmacological properties. International Journal of Molecular Sciences. 2021 Jan 9;22(2):618. 10.3390/ijms22020618

11) Saha P, Sharma D, Dash S, Dey KS, Sil SK. Identification of 2, 4-Di-tert-butylphenol (2, 4-DTBP) as the Major Contributor of Anti-colon cancer Activity of Active Chromatographic Fraction of Parkia javanica (Lamk.) Merr. Bark Extract. Biomedical and Pharmacology Journal. 2023 Mar 21;16(1):275–88.

12) Farahpour MR, Mirzakhani N, Doostmohammadi J, Ebrahimzadeh M. Hydroethanolic Pistacia atlantica hulls extract improved wound healing process; evidence for mast cells infiltration, angiogenesis and RNA stability. International Journal of Surgery. 2015 May 1;17:88–98. 10.1016/j.ijsu.2015.03.019

13) Farahpour MR, Sheikh S, Kafshdooz E, Sonboli A. Accelerative effect of topical Zataria multiflora essential oil against infected wound model by modulating inflammation, angiogenesis, and collagen biosynthesis. Pharmaceutical Biology. 2021 Jan 1;59(1):1–0. 10.1080/13880209.2020.1861029

14) SK HZ, Farahpour MR, Kar HH. The Effect of Topical Administration of an Ointment Prepared From Trifolium repens Hydroethanolic Extract on the Acceleration of Excisional Cutaneous Wound Healing. Wounds: a Compendium of Clinical Research and Practice. 2020 Sep 1;32(9):253–61. 10.25270/wnds/2020.253261

15) Dardmah F, Farahpour MR. Quercus infectoria gall extract aids wound healing in a streptozocin-induced diabetic mouse model. Journal of Wound Care. 2021 Aug 2;30(8):618–25. 10.12968/jowc.2021.30.8.618

16) Farahpour MR, Pirkhezr E, Ashrafian A, Sonboli A. Accelerated healing by topical administration of Salvia officinalis essential oil on Pseudomonas aeruginosa and Staphylococcus aureus infected wound model. Biomedicine & Pharmacotherapy. 2020 Aug 1;128:110120. 10.1016/j.biopha.2020.110120

17) Bickford PC, Tan J, Shytle RD, Sanberg CD, El-Badri N, Sanberg PR. Nutraceuticals synergistically promote proliferation of human stem cells. Stem cells and development. 2006 Feb 1;15(1):118–23. 10.1089/scd.2006.15.1

18) Shen CL, Kwun IS, Wang S, Mo H, Chen L, Jenkins M, Brackee G, Chen CH, Chyu MC. Functions and mechanisms of green tea catechins in regulating bone remodeling. Current Drug Targets. 2013 Dec 1;14(13):1619–30. https://10.2174/13894501113146660216

19) Jaźwińska A, Sallin P. Regeneration versus scarring in vertebrate appendages and heart. The Journal of pathology. 2016 Jan;238(2):233–46. 10.1002/path.4644

20) Gemberling M, Bailey TJ, Hyde DR, Poss KD. The zebrafish as a model for complex tissue regeneration. Trends in genetics. 2013 Nov 1;29(11):611–20. 10.1016/j.tig.2013.07.003

21) Tavares B, Lopes SS. The importance of Zebrafish in biomedical research. Acta medica portuguesa. 2013 Oct 31;26(5):583–92. 10.20344/amp.4628

22) Xu C, Volkery S, Siekmann AF. Intubation-based anesthesia for long-term time-lapse imaging of adult zebrafish. Nature Protocols. 2015 Dec;10(12):2064–73. 10.1038/nprot.2015.130

23) Teixidó E, Piqué E, Gómez-Catalán J, Llobet JM. Assessment of developmental delay in the zebrafish embryo teratogenicity assay. Toxicology in Vitro. 2013 Feb 1;27(1):469–78. 10.1016/j.tiv.2012.07.010

24) Jaźwińska A, Badakov R, Keating MT. Activin-βA signaling is required for zebrafish fin regeneration. Current biology. 2007 Aug 21;17(16):1390–5. DOI 10.1016/j.cub.2007.07.019

25) Stoick-Cooper CL, Weidinger G, Riehle KJ, Hubbert C, Major MB, Fausto N, Moon RT. Distinct Wnt signaling pathways have opposing roles in appendage regeneration. 10.1242/dev.001123

26) Whitehead GG, Makino S, Lien CL, Keating MT. fgf20 is essential for initiating zebrafish fin regeneration. Science. 2005 Dec 23;310(5756):1957–60. DOI: 10.1126/science.1117637

27) Chablais F, Jaźwińska A. IGF signaling between blastema and wound epidermis is required for fin regeneration. Development. 2010 Mar 15;137(6):871–9. 10.1242/dev.043885

28) Blum N, Begemann G. Retinoic acid signaling controls the formation, proliferation and survival of the blastema during adult zebrafish fin regeneration. Development. 2012 Jan 1;139(1):107–16. 10.1242/dev.065391

29) Dhakal R, Kalladka K, Singha A, Pandyanda Nanjappa D, Ravindra J, Vittal R, Sil SK, Chakraborty A, Chakraborty G. Investigation of anti-proliferative and anti-angiogenic properties of Parkia javanica bark and fruit extracts in zebrafish. Plos one. 2023 Jul 21;18(7):e0289117. 10.1371/journal.pone.0289117

30) Zainol Abidin IZ, Fazry S, Jamar NH, Ediwar Dyari HR, Zainal Ariffin Z, Johari AN, Ashaari NS, Johari NA, Megat Abdul Wahab R, Zainal Ariffin SH. The effects of Piper sarmentosum aqueous extracts on zebrafish (Danio rerio) embryos and caudal fin tissue regeneration. Scientific reports. 2020 Aug 25;10(1):14165. 10.1038/s41598-020-70962-7

31) Chen F, Pu S, Tian L, Zhang H, Zhou H, Yan Y, Hu X, Wu Q, Chen X, Cheng SH, Xu S. Radix Rehmanniae Praeparata promoted zebrafish fin regeneration through aryl hydrocarbon receptor-dependent autophagy. Journal of ethnopharmacology. 2024 Sep 15;331:118272. 10.1016/j.jep.2024.118272

32) Zhang P, Liu N, Xue M, Zhang M, Xiao Z, Xu C, Fan Y, Liu W, Qiu J, Zhang Q, Zhou Y. Antiinflammatory and antioxidant properties of squalene in copper sulfate-induced inflammation in zebrafish (Danio rerio). International journal of molecular sciences. 2023 May 10;24(10):8518. 10.3390/ijms24108518

33) Sarkar A, Chakrabarti A, Bhaumik S, Debnath B, Singh SS, Ghosh R, Zaki ME, Al-Hussain SA, Debnath S. Parkia javanica edible pods reveal potential as an anti-diabetic agent: UHPLC-QTOF-MS/MS-Based chemical profiling, in silico, in vitro, in vivo, and oxidative stress studies. Pharmaceuticals. 2024 Jul 21;17(7):968. doi:10.3390/ph17070968

34) Saleh MS, Jalil J, Zainalabidin S, Asmadi AY, Mustafa NH, Kamisah Y. Genus Parkia: Phytochemical, medicinal uses, and pharmacological properties. International Journal of Molecular Sciences. 2021 Jan 9;22(2):618. 10.3390/ijms22020618

35) Banu S, Gaur N, Nair S, Ravikrishnan T, Khan S, Mani S, Bharathi S, Mandal K, Kuram NA, Vuppaladadium S, Ravi R. Understanding the complexity of epimorphic regeneration in zebrafish caudal fin tissue: A transcriptomic and proteomic approach. Genomics. 2022 Mar 1;114(2):110300. 10.1016/j.ygeno.2022.110300

36) Cao Z, Yang Q, Luo L. Zebrafish as a model for germ cell regeneration. Frontiers in Cell and Developmental Biology. 2021 Jul 22;9:685001. doi:10.3389/fcell.2021.685001

37) Wang G, Yang F, Zhou W, Xiao N, Luo M, Tang Z. The initiation of oxidative stress and therapeutic strategies in wound healing. Biomedicine & Pharmacotherapy. 2023 Jan 1;157:114004. 10.1016/j.biopha.2022.114004

38) Schieber M, Chandel NS. ROS function in redox signaling and oxidative stress. Current biology. 2014 May 19;24(10):R453–62. DOI: 10.1016/j.cub.2014.03.034

39) Gupta A, Singh RL, Raghubir R. Antioxidant status during cutaneous wound healing in immunocompromised rats. Molecular and Cellular Biochemistry. 2002 Dec;241(1):1–7. DOI: 10.1023/a:1020804916733

40) Rajiv C, Roy SS, Tamreihao K, Kshetri P, Singh TS, Sanjita Devi H, Sharma SK, Ansari MA, Devi ED, Devi AK, Langamba P. Anticarcinogenic and antioxidant action of an edible aquatic flora jussiaea repens L. Using in vitro bioassays and in vivo zebrafish model. Molecules. 2021 Apr 15;26(8):2291. 10.3390/molecules26082291

41) Rajiv C, Devi HS, Devi AK, Tamreihao K, Kshetri P, Tania C, Singh TS, Sonia C, Singh MN, Sen A, Sharma SK. Pharmacological potential of Jussiaea repens L. against CuSO4 and bacterial lipopolysaccharide O55: B5 induced inflammation using in-vivo zebrafish models. Journal of Ethnopharmacology. 2024 Jan 10;318:116932. 10.1016/j.jep.2023.116932

42) Solarte David VA, Güiza-Argüello VR, Arango-Rodríguez ML, Sossa CL, Becerra-Bayona SM. Decellularized tissues for wound healing: towards closing the gap between scaffold design and effective extracellular matrix remodeling. Frontiers in bioengineering and biotechnology. 2022 Feb 16;10:821852. 10.3389/fbioe.2022.821852

43) Saxena S, Purushothaman S, Meghah V, Bhatti B, Poruri A, Meena Lakshmi MG, Sarath Babu N, Narasimha Murthy CL, Mandal KK, Kumar A, Idris MM. Role of annexin gene and its regulation during zebrafish caudal fin regeneration. Wound repair and regeneration. 2016 May;24(3):551–9. DOI: 10.1111/wrr.12429

44) Walczyńska KS, Zhu L, Liang Y. Insights into the role of the Wnt signaling pathway in the regeneration of animal model systems. The International journal of developmental biology. 2023 Oct 31;67(3):65–78. 10.1387/ijdb.220144yl

